# Characterization of a novel *Fgf10^CreERT2^* knock-in mouse line targeting postnatal lung Fgf10 lineages

**DOI:** 10.1101/2021.02.05.429562

**Authors:** Xuran Chu, Sara Taghizadeh, Ana Ivonne Vasquez-Armendariz, Susanne Herold, Lei Chong, Chengshui Chen, Jin-San Zhang, Elie El Agha, Saverio Bellusci

## Abstract

*Fgf10* is a key gene during development, homeostasis and repair after injury. We previously reported a *Fgf10^CreERT2^* line (with the CreERT2 cassette inserted in frame with the start codon of exon 1), called thereafter *Fgf10^Ki-v1^*, to target Fgf10^Pos^ cells. While this line allowed fairly efficient and specific labeling of Fgf10^Pos^ cells during the embryonic stage, it failed to target these cells after birth, particularly in the postnatal lung, which has been the focus on our research. We report here the generation and validation of a new *Fgf10^CreERT2^* (called thereafter *Fgf10^Ki-v2^*) with the insertion of the expression cassette in frame with the stop codon of exon 3. This new *Fgf10^Ki-v2^* line exhibited comparable *Fgf10* expression level *to their wild type counterpart*. However, a disconnection between the *Fgf10* and the *Cre* expression was observed in *Fgf10^Ki-v2/+^* lungs. In addition, lung and limb agenesis were observed in homozygous embryos suggesting a loss of *Fgf10* functional allele in *Fgf10^Ki-v2^* mice. Bio-informatics analysis shows that the 3’UTR, where the CreERT2 cassette is inserted, contains numerous putative transcription factor binding sites. By crossing this line with tdTomato reporter line, we demonstrated that tdTomato expression faithfully recapitulated Fgf10 expression during development. Significantly, *Fgf10^Ki-v2^* mouse is capable of significantly targeting Fgf10^Pos^ cells in the adult lung. Therefore, despite the aforementioned limitations, this new *Fgf10^Ki-v2^* line opens the way for future mechanistic experiments involving the postnatal lung.

## Introduction

The Fibroblast growth factor (FGF) family consisting of 22 members is divided into three groups: the paracrine FGF group signaling through FGFR and heparin-sulfate proteoglycans, the endocrine FGF group signaling through FGFR with Klotho family of proteins as co-receptors, and the intracellular FGF group involved in FGFR independent signaling [1]. The FGF7 subgroup which contains FGF3, 7, 10, 22 belongs to the paracrine FGF group. These growth factors interact mostly with the FGFR2b receptor. FGF10 in particular has been shown to play important roles during development, homeostasis and repair after injury. In the lung, it plays a crucial role in regulating the branching morphogenesis [2]. Genetic deletion of either *Fgf10* or its predominant receptor *Fgf2b* leads to agenesis of both the limb and the lung, specific portions of the gut, the pancreas as well as the mammary, lacrimal and salivary glands. During homeostasis, Fgfr2b signaling has been shown to be critical for the regeneration of the incisors in mice as well as for the maintenance of the terminal end buds in the mammary gland [3]. In the context of the repair process, *Fgf10* deletion in peribronchial mesenchymal cells leads to impaired repair following bronchial epithelium injury using naphthalene [4, 5]. On the other hand, over-expression of *Fgf10* reduces the severity of lung fibrosis in bleomycin-induced mice. Given these diverse biological activities, it is important to generate and validate mouse knock in lines to monitor the localization, fate and status of Fgf10^Pos^ cells during development, homeostasis and repair after injury.

We have previously generated a *Fgf10^Cre-ERT2^* knock in mice line, called thereafter *Fgf10^KI-v1^* mice, to monitor the fate of Fgf10^Pos^ cells after tamoxifen (Tam) administration [6]. In these mice, the Tam-inducible Cre recombinase (Cre-ERT2-IRES-YFP) was inserted in frame with the start codon of the endogenous *Fgf10* gene. *Fgf10^KI-v1^* corresponds to a loss-of-function allele for *Fgf10* as evidenced by our observation that *Fgf10^KI-v1/KI-v1^* homozygous embryos die at birth from multiple organ agenesis, including the lung. In the *Fgf10^KI-v1/+^* lungs, the expression of *Cre* recombinase gradually decreases to almost undetectable levels postnatally, rendering impossible the monitoring of Fgf10^Pos^ cells postnatally. This is likely due to the deletion of intronic sequences containing key transcription binding sites at the insertion of the Cre-expression cassette.

In order to circumvent this problem, we therefore generated a new Cre-ERT2 knock in line (named *Fgf10^KI-v2^*) by targeting the 3’UTR of the *Fgf10* gene. We here provide experimental evidence for the validation of these mice. Besides a PCR based strategy to genotype the *Fgf10^KI-v2^* allele, we have also established a qPCR-based approach to monitor the expression level of *Fgf10* and *Cre* at different developmental stages in the lung. *Fgf10^KI-v2/KI-v2^* homozygous embryos have been generated to check for developmental defects. *Fgf10^KI-v2/+^* lines were crossed with the *tdTomato^flox^* mice to validate, at two different embryonic stages, the expression patterns of tdTomato in previously known domains of *Fgf10* expression. Importantly, we validated the use of these mice in the adult stages to target Fgf10^Pos^ cells in the lung. Flow cytometry analysis and immunofluorescence staining has been carried out to characterize further these cells along the lipofibroblast lineage. Bio-informatic analysis of the insertion site of the CreERT2 cassette in the 3’UTR has been carried out. Altogether, our results indicate that we have generated successfully a new *Fgf10^CreERT2^* line to target Fgf10^Pos^ cells both in embryonic and adult stages.

## Material and methods

### Genotyping

Two pairs of primers were used to determine the genotype of *Fgf10^Ki-v2^* knock in mice line. Primer P1 (5’-AACACCTCTGCTCACTTCCTC-3’); and primer P2 (5’-AGGGTCCACCTTCCGCTTTT-3’) were used to detect the knock-in allele (251bp band) whereas primer P3 (5’-GCAGGCAAATGTATGTGGCA-3’) and primer P4 (5’-TGCTTGCGTGTCTTACTGCT-3’) were used to detect the wild-type allele (580bp band). The PCR program consists of a denaturation step at 94°C for 2 min, followed by 35 cycles of denaturation (94 °C for 30s), annealing (65 °C for 30s) and extension steps (68°C for 300s). The program ends with a completion step at 68°C for 480s. Each PCR tube contains 4.3 μl of H_2_O, Taq DNA Polymerase in 5.5 μL of Qiagen Master Mix (QIAGEN, Hilden, Germany), 15 pmol of each primer, and 10 ng of genomic DNA in a final volume of 11 μL.

### Mice and Tamoxifen Administration

All mice were kept under specific pathogen free (SPF) conditions with unlimited food and water. *tdTomato^flox/flox^* reporter mice were purchased from Jackson lab (B6; 129S6-Gt(ROSA)^26Sortm9(CAG-tdTomato)Hze^/J, ref 007905). Embryonic day 0.5 (E0.5) was assigned to the day when a vaginal plug was detected. Mice were housed in an SPF environment. Animal experiments proposals were approved by the Regierungspraesidium Giessen (approval number RP GI/47-2019). Tamoxifen stock solution was prepared by dissolving tamoxifen powder (Sigma, T5648-5G) in corn oil at a concentration of 20 mg/mL at room temperature and stored in −20°C. Adult mice received 3 successive Tam-IP (0.075 mg tamoxifen per gram of body weight) before sacrificed. Pregnant female mice and pups received one intra-peritioneal [7] injection. Dissected mice were examined using Leica M205 FA fluorescent stereoscope (Leica, Wetzlar, Germany) and images were acquired using Leica DFC360 FX camera. Figures were assembled in Adobe Photoshop CS5.

### Quantitative Real-time PCR and Statistical Analysis

Freshly isolated embryos and lungs were lysed and RNA was extracted using RNeasy kit (Qiagen, Hilden, Germany). Freshly dissected lungs of E13.5 embryos were homogenized using QiaShredder columns (Qiagen). 1000 ng of RNA was used for cDNA synthesis using Quantitect Reverse Transcription kit (Qiagen). Primers and probes for *Fgf10*, *Cre* and *B2M* were designed using NCBI Primer-BLAST (https://www.ncbi.nlm.nih.gov/tools/primer-blast/). More details about the used primers and probes can be found in Table S1. Quantitative real-time PCR (qPCR) was performed using LightCycler 480 real-time PCR machine (Roche Applied Science). Samples were run in doublets using B2M as a reference gene and the DDCT method was used to calculate the relative quantification. GraphPad Prism software was used to generate and analysis data. statistical analyses were performed using Student’s t-test (for comparing two groups) or One-way ANOVA (for comparing three or more groups). Data were considered significant if P<0.05.

### Flow cytometry

Freshly dissected lung washed with Hanks’ balanced salt solution (HBSS) and kept on ice. Then using sharp blades to cut the lung into small pieces and digested with 0.5% collagenase Type IV in HBSS (Life Technologies, Invitrogen) for 45 minutes at 37 °C. Passing the lung homogenates through 18, 21 and 24G syringes followed by passing through 70 and 40 cell strainers (BD Biosciences, Heidelberg, Germany). Centrifuge lung homogenates at 1000 rpm at 4 °C for 10 minutes and then re-suspend cell pellets in HBSS. 20 μl of sample was taken as an unstained control. Apply antibodies (Biolegend, Fell, Germany) against Cd45 (APC-conjugated; 1:50), Cd31 (APC-conjugated; 1:50), Epcam (APC-Cy7-conjugated; 1:50) and Sca1 (Pacific blue-conjugated; 1:50) as well as LipidTOX staining (FITC-conjugated, 1:200) for 30 minutes on ice in the. Add 1 ml HBSS and then centrifuge at 4 °C for 5 minutes to wash. Then re-suspended the cell pellet in HBSS. FACSAria III cell sorter (BD Biosciences) was used to carry out the FACs measurements and sorting. Endogenous tdTomato signal was detected through PE channel. Gates were set up according to the unstained controls

### Bio-informatics

NCBI genes was used to find the mouse murine *Fgf10* sequence, the identification of putative transcription factor binding site (TFBS) was done using the online PROMO software (http://alggen.lsi.upc.edu/recerca/menu_recerca.html). The list of putative TFBS located on area including the exon 3 and the 3’UTR were further compared to the previously identified transcription factors expressed in the lung mesenchyme [8].

## Results

### Generation of a novel *Fgf10* knock in line (*C57BL6-Fgf10^tm2(YFP-CreERT2)Sbel^/J* aka *Fgf10^KI-v2^*) with the insertion of the Cre-ERT2 cassette in frame with the stop codon of Exon3

129Sv ES cells were electroporated with a targeting vector containing the first 3kb of exon 3 of the *Fgf10* open reading frame (Fig.1A, B). Immediately downstream of the stop codon of exon 3 is the F2A sequence encoding for the self-cleaving peptide, followed by the coding sequence of eYFP, the self-cleaving peptide sequence T2A, the tamoxifen-inducible form of Cre recombinase (Cre-ERT2) [17], and the Neomycin-resistance gene (Neo) respectively. Resistant ES cell clones were selected, screened by PCR and then verified by Southern blotting. Selected ES clones were injected into C57BL/6J blastocysts to generate chimeric pups (Fig. 1C). Chimeras were then crossed with C57BL/6J mice ubiquitously expressing Flp recombinase to generate heterozygous *Fgf10^Ki-v2^* knock-in mice where the Neo cassette was totally excised (Fig. 1D). Genotyping strategy with primers P1/P2 (with P1 located just before the STOP codon and P2 being part of the F2A sequence) to detect the *Fgf10^Ki-v2^* mutant allele and P3/P4 (located before and after the STOP codon in exon 3, respectively) to detect the *Fgf10^+^* wild type allele (Fig. 1E).

**Figure 1.**
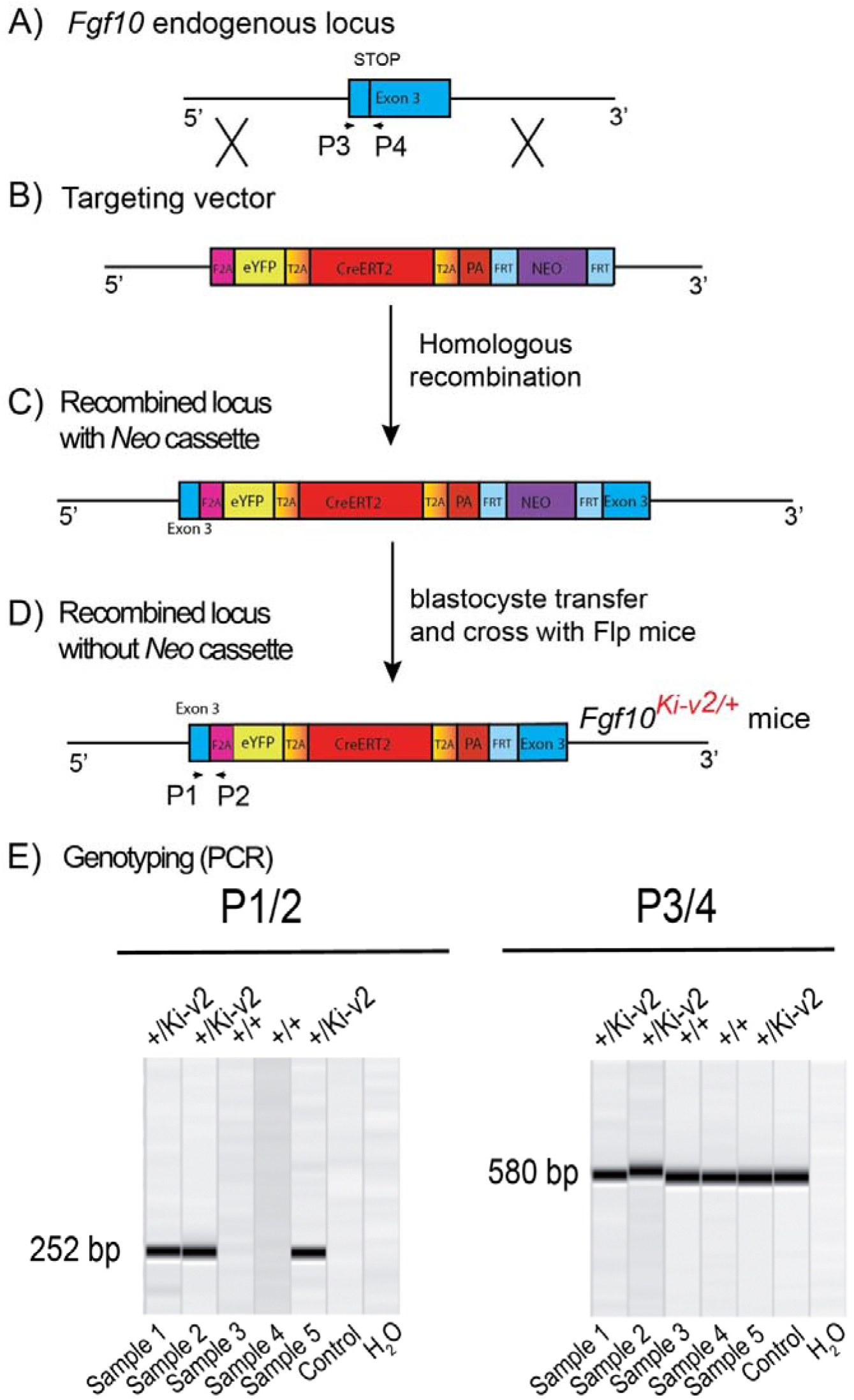
Generation and genotyping of the novel *Fgf10^Ki-v2^* line. **(A, B)** Homologous recombination was carried out to insert the F2A-eYFP-T2A-CreERT2-T2A-PA-NEO construct in frame with the stop codon of exon 3 of the mouse *Fgf10* gene. Neomycin resistance coding gene was used for the positive selection. **(C, D)** Recombined ES cells clones were treated with flipase to remove the Neo cassette and blastocyste transfer of the selected ES cells was carried out to generate chimera animals. **(E)** PCR strategy to genotype mutant and wild type animals. Primers 1 and 2 were used for the detection of the mutant *Fgf10^Ki-v2^* allele (252bp) and Primers 3 and 4 were used for the detection of the wild type *Fgf10^+^* allele (580bp).

### The *Fgf10^Ki-v2^* is a loss of function allele

Our initial design of the novel *Fgf10^Ki-v2^* knock in line targeting the 3’UTR was conceived to allow normal expression of *Fgf10*. We carried out the initial validation for *Fgf10* expression in *Fgf10^Ki-v2/+^* vs. *Fgf10^+/+^* (WT) in the lung of embryonic and postnatal mice isolated at different time-points (Fig. 2A). Our results indicated that *Fgf10* expression level in *Fgf10^Ki-v2/+^* lungs is comparable to the one observed in the *Fgf10^+/+^* lungs at all these time-points (Fig. 2B). Next, we compared *Fgf10* vs. *Cre* expression in *Fgf10^Ki-v2/+^* lungs at different time-points (Fig. 2C). Our results indicate a lower level of *Cre* compared to *Fgf10* at all these time-points (Fig. 2D). This, disconnect between *Cre* and *Fgf10* expression suggests that the insertion of the CreERT2 cassette in the 3’UTR disrupted the expression of the endogenous *Fgf10* gene produced from the recombined allele. To determine whether the insertion of Cre-ERT2 in the endogenous *Fgf10* locus led to loss of function of *Fgf10*, *Fgf10^Ki-v2^* heterozygous animals were crossed together and embryos were harvested at E13.5. *Fgf10^Ki-v2/Ki-v2^* homozygous embryos suffered from lung and limb agenesis, which is consistent with complete loss of function of *Fgf10* (Fig. 2E). qPCR revealed minimal expression levels for *Fgf10* in *Fgf10^Kii-v2/Ki-v2^* embryos (n = 3) compared to *Fgf10^Ki-v2/+^(*n=3; P=0.0001) and *Fgf10^+/+^* embryos (n= 3; P=0.0001) (data not shown).

**Figure 2.**
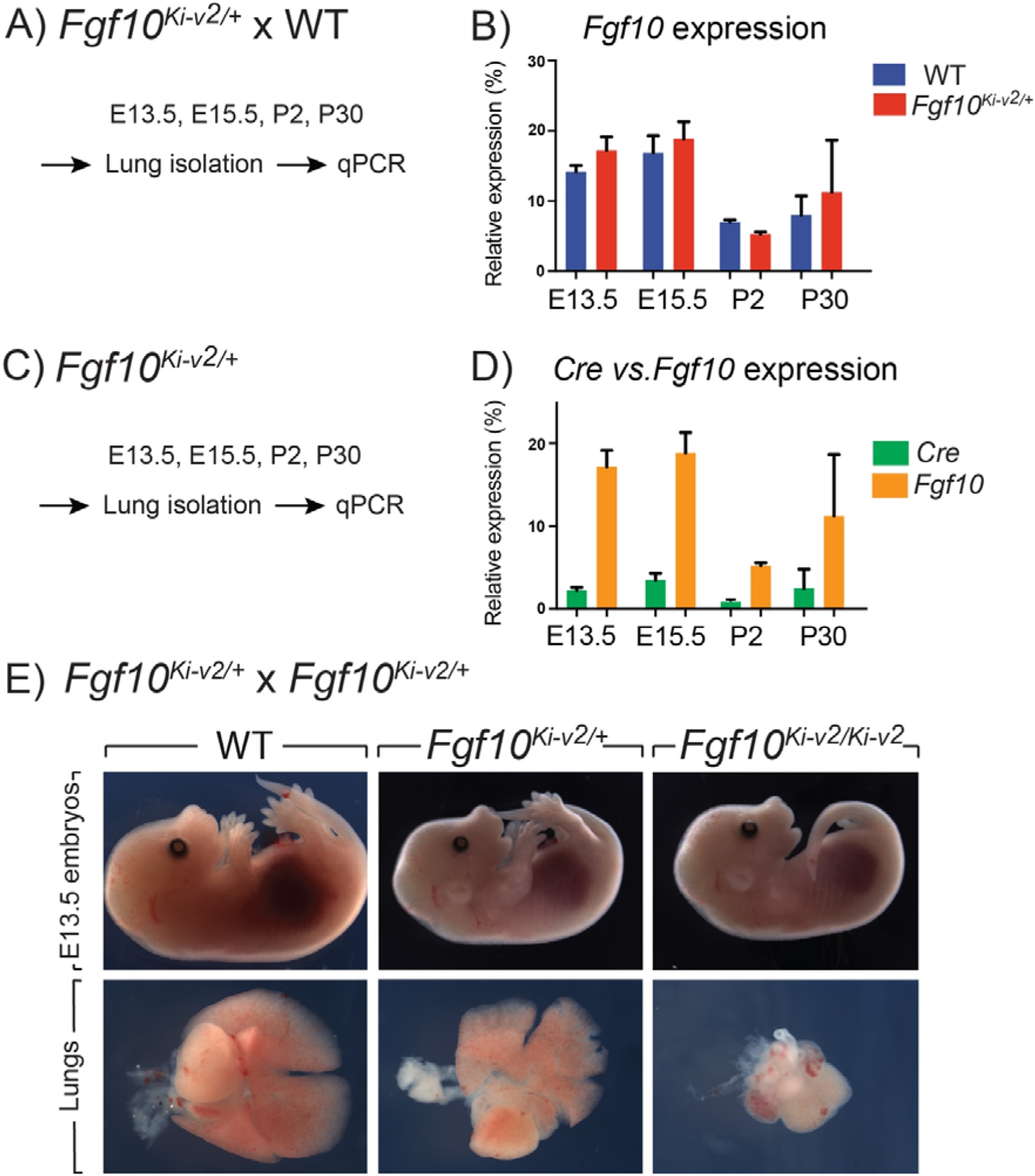
*Fgf10^Ki-v2^* is a loss of function allele. **(A)** *Fgf10^Ki-v2/+^* and *Fgf10^+/+^* mice were crossed to generate animals at different staged. The lungs were isolated at E13.5, E15.5, P2 and P30 and processed for qPCR. **(B)** qPCR results for *Fgf10* expression in *Fgf10^+/+^* and *Fgf10^Ki-v2/+^* at these time points. **(C, D)** Generation of *Fgf10^Ki-v2/+^* animals at different time points. The lungs were isolated at E13.5, E15.5, P2 and P30 and processed for qPCR to monitor *Cre* and *Fgf10* expression. **(E)** *Fgf10^Ki-v2/+^* males and females were crossed to generate *Fgf10^+/+^*, *Fgf10^Ki-v2/+^* and *Fgf10^Ki-v2/Ki-v2^* embryos at E13.5. Note the absence of limbs and lungs in the *Fgf10^Ki-v2/Ki-v2^* embryos.

We therefore conclude that the *Fgf10^Ki-v2^* allele corresponds to a *Fgf10* loss of function allele. This also indicates that *Fgf10* expression in *Fgf10^Ki-v2/+^* heterozygous lungs arises mostly from the non-recombined *Fgf10* allele.

### Validation of Cre activity to label Fgf10-positive Cells during embryonic development

In order to test the recombinase activity of Cre-ERT2, *Fgf10^Ki-v2/+^* heterozygous mice were crossed with *tdTomato^flox/flox^* reporter mice. Pregnant female mice received a single IP injection of tamoxifen at E11.5 (Fig. 3A) or E15.5 (Fig. 4A). Embryos were harvested at E18.5. No fluorescent signal was observed in *Fgf10^+/+^; tdTomato^flox/+^* embryos (Fig. 3B and Fig. 4B; n=4) indicating absence of recombination in control embryos and lack of leakiness of the *tdTomato^flox^* allele. By contrast, tamoxifen treatment at E11.5 led to a strong fluorescent signal in the limbs, stomach, cecum, colon and lungs of *Fgf10^Ki-v2/+^;Tomato^flox/+^* embryos (n=3). In the limb, the labeled cells were more abundant in the digit tip area, known to express high level of *Fgf10* [9]. Along the gastro-intestinal tract, labeled cells were located in the anterior part of the stomach as well as in duodenum (data not shown) which are both reported to express high level of *Fgf10* [10]. A similar observation was made in the cecum and the distal colon [10]. Throughout the lung, we found a robust tdTomato expression with a higher expression in the interlobular septa. This is similar to what was observed with the previously validated *Fgf10^LacZ^* reporter line [11]. Interestingly, in the trachea, no labeled cells were observed in this experimental condition (Fig. 3C). Additionally, tamoxifen treatment at E15.5 revealed strong fluorescent signal in the pinna of the developing ear as well as in the trachea and in between the cartilage rings. These two additional expression domains are consistent with sites of *Fgf10* expression [12, 13]. We therefore conclude that Cre expression reflects *Fgf10* expression and that this line can be used to target Fgf10^Pos^ cells.

**Figure 3.**
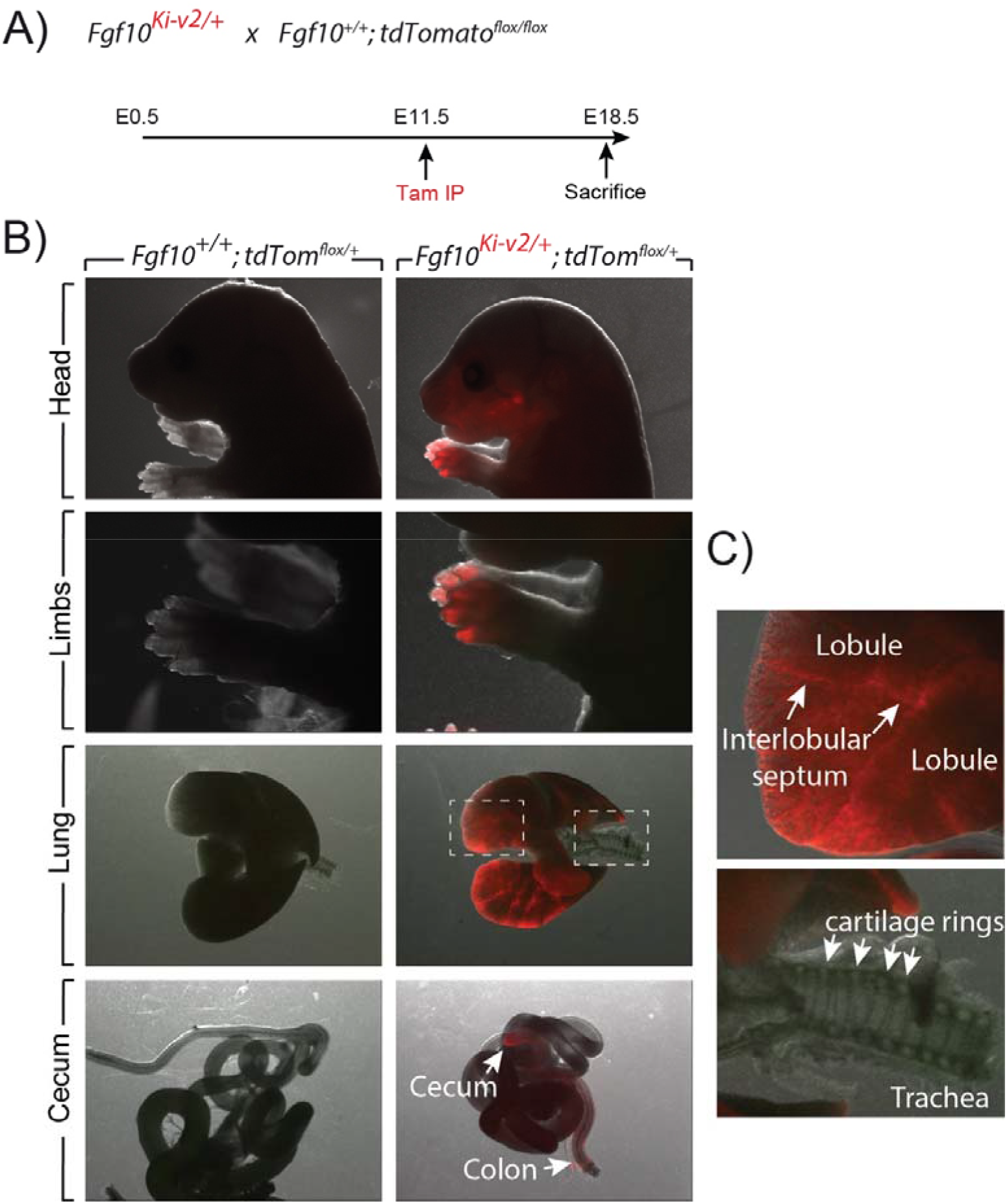
Validation of the labeling of Fgf10^Pos^ cells at E11.5. **(A)** *Fgf10^Ki-v2/+^* were crossed with *Fgf10^+/+^*; *tdTom^flox/flox^* mice. Pregnant females received a single injection of tamoxifen IP when the embryos were at E11.5 and sacrificed at E18.5. **(B)** Head, limbs, lung and cecum of *Fgf10^+/+^*; *tdTom^flox/+^* and *Fgf10^Ki-v2/+^*; *tdTom^flox/+^* embryos are shown. Note the absence of fluorescence in the *Fgf10^+/+^; tdTom^flox/+^* indicating that the non-recombined *LoxP-Stop-LoxP* tdTomato allele is not leaky. **(C)** Higher magnification of lung and trachea showing enriched tdTomato expression in the interlobular septa and the lack of tdTomato expression between the cartilage rings, respectively.

**Figure 4.**
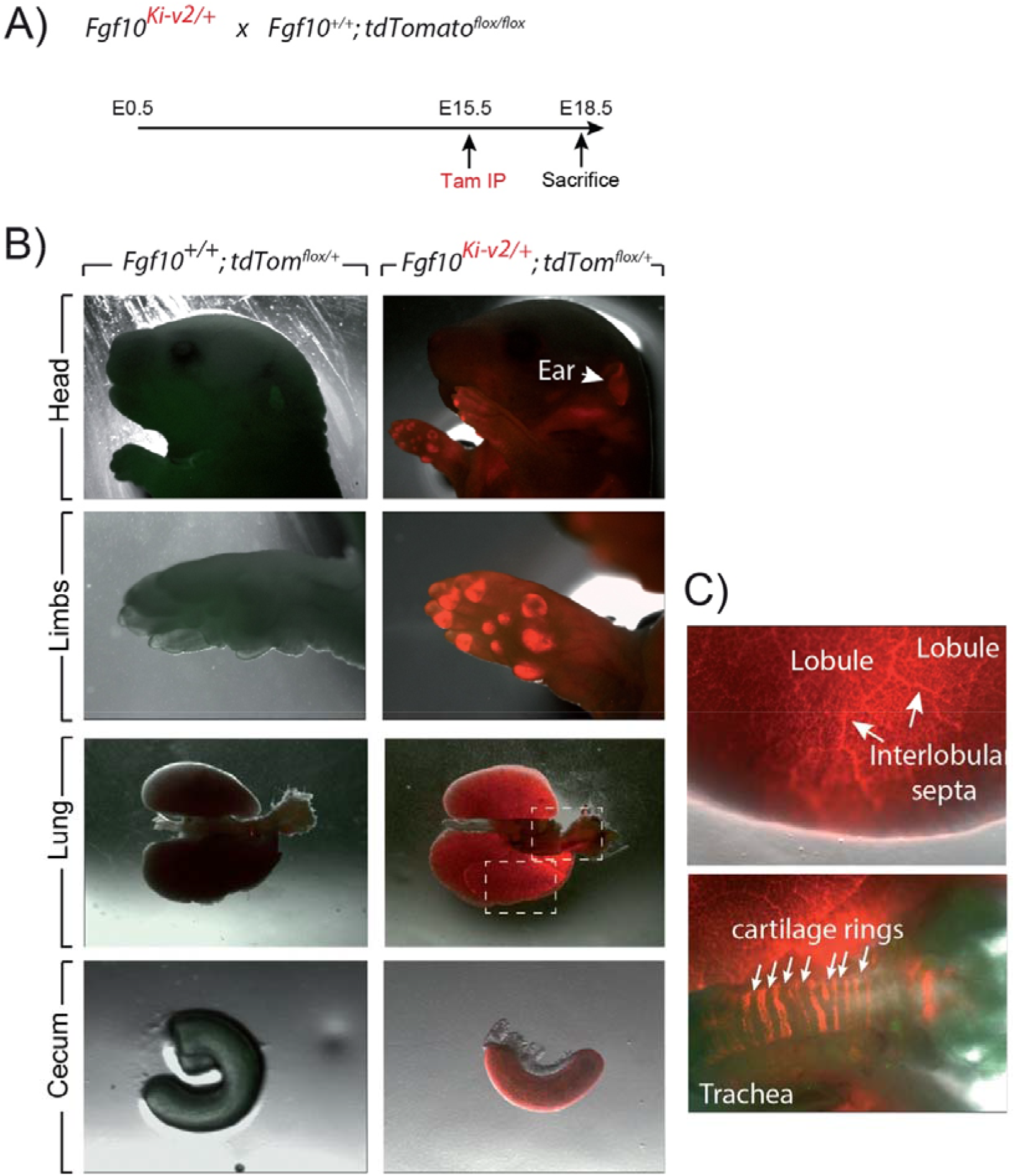
Validation of the labeling of Fgf10^Pos^ cells at E15.5. **(A)** *Fgf10^Ki-v2/+^* were crossed with *Fgf10^+/+^*; *tdTom^flox/flox^* mice. Pregnant females received a single injection of tamoxifen IP when the embryos were at E15.5 and sacrificed at E18.5. **(B)** Head, limbs, lung and cecum of *Fgf10^+/+^*; *tdTom^flox/+^* and *Fgf10^Ki-v2/+^*; *tdTom^flox/+^* embryos are shown. Note the expression in the external ear (arrow) **(C)** Higher magnification of lung and trachea showing enriched tdTomato expression in the interlobular septa and the presence of tdTomato expression between the cartilage rings, respectively.

### Fgf10^Pos^ cells labeled after birth do not contribute significantly to the secondary crest myofibroblast during alveologenesis

Using the previously generated Fgf10^Ki-v1/+^ line, we demonstrated that Fgf10^Pos^ cells do not contribute in a major way to the Acta2^Pos^ secondary crest myofibroblasts (SCMF) which are abundant during the first 2-3 weeks during alveologenesis which takes place from postnatal day 5 (P5) to -P28 (El Agha et al., 2014). To confirm this observation with the new Fgf10^Ki-v2/+^ line, we captured Fgf10^Pos^ cells at P4 and examined the status of the labeled cells at P21, one week before the end of the alveologenesis phase (Fig. 5A). Analysis of the whole lung by fluorescence videomicroscopy indicated a much higher number of labeled cells in the *Fgf10^Ki-v2^; tdTomato^flox/+^* lung compared to the *Fgf10^Ki-v1;^ tdTomato^flox/+^* lung (Figure 5B). IF for Acta2 on these lungs was also carried out and quantified in the two lines (Fig. 5C and D). We also used previously generated PFA-fixed thin lung sections for the *Fgf10^Ki-v1;^ tdTomato^flox/+^* lung. Our results indicate that that a higher % of Tom^Pos^/DAPI is observed on sections of *Fgf10^Ki-v2^; tdTomato^flox/+^* vs. *Fgf10^Ki-v1;^ tdTomato^flox/+^* (4.7% ± 0.6% vs. 1.5% ± 0.2%, n=2) thereby confirming the fluorescence videomicroscopy results (Fig. 5B). Interestingly, the percentile of Tom^Pos^Acta2^Pos^/Tom^Pos^ is much reduced in the *Fgf10^Ki-v2^; tdTomato^flox/+^* vs. *Fgf10^Ki-v1;^ tdTomato^flox/+^* lungs (4.7% ± 0.9% vs. 25.3% ± 5.9%, n=2) indicating minimal commitment of the Fgf10^Pos^ cells to the SCMF lineage. This difference between the two lines could be explained by the relatively low number of cells present in the *Fgf10^Ki-v1;^ tdTomato^flox/+^* lungs. In addition, while we used thick vribratome sections and confocal to quantify the cells in *Fgf10^Ki-v2^; tdTomato^flox/+^* lungs, we used previously generated IF images on PFA fixed sections for *Fgf10^Ki-v1;^ tdTomato^flox/+^* lungs. The higher number of cells counted and the higher resolution by confocal microscopy due to the use of thick sections could therefore explain this difference. Interestingly, the quantification of Tom^Pos^Acta2^Pos^/Acta2^Pos^ indicated no significant difference between *Fgf10^Ki-v2^; tdTomato^flox/+^* and *Fgf10^Ki-v1;^ tdTomato^flox/+^* lungs suggesting overall a similar a relatively low contribution of the Fgf10^Pos^ cells to the SCMF lineage (5.2% ± 1.7% vs. 3.1% ± 0.7%, n=2)

**Figure 5.**
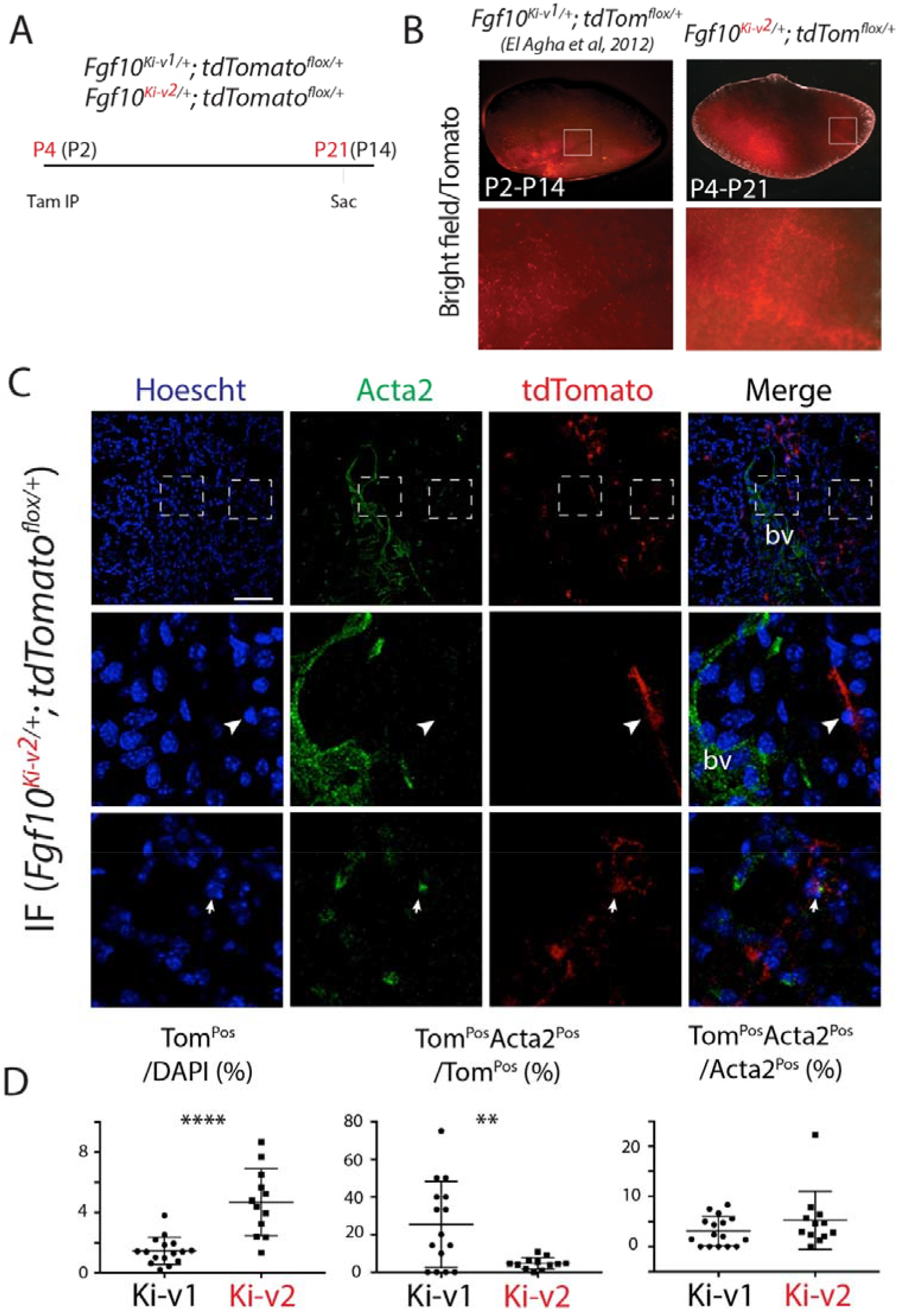
Fgf10^Pos^ cells labeled after birth do not contribute significantly to the secondary crest myofibroblast during alveologenesis. **(A)** Fgf10^Pos^ cells in *Fgf10^Ki-v2/+^; tdTomato^flox/+^* pups were labeled in vivo at P4 and analyzed at P21. We also used previously generated *Fgf10^Ki-v1/+^; tdTomato^flox/+^* samples induced at P2 and analyzed at P14 **(B)** Whole-mount fluorescent images of *Fgf10^Ki-v2/+^*; *tdTom^flox/+^* and *Fgf10^Ki-v1/+^*; *tdTom^flox/+^* lungs showing more abundant labeled cells in *Fgf10^Ki-v2/+^* vs. *Fgf10^Ki-v1/+^* lungs. (**C**) Acta2 IF on *Fgf10^Ki-v2/+^*; *tdTom^flox/+^*shows little contribution of Fgf10^Pos^ cells to the SCMF (Acta2^Pos^Tom^Pos^/Acta2^Pos^). **(D**) Quantification of Tom^Pos^ cells.

### The new *Fgf10^Ki-v2^* line allows enhanced labeling of Fgf10^Pos^ cells in the adult lung compared to the previous *Fgf10^KI-v1^* line

2 months-old *Fgf10^Ki-v1/^*^+^; *tdTomato^flox/flox^* and *Fgf10^K-v2^*^/+^; *tdTomato^flox/flox^* mice and were treated with Tam IP or oil at day 1 (D61), 3 (D63) and 5(D65) and the lungs were collected at day 7 (D67) (Fig. 6A). No fluorescent signal was observed in oil-treated *Fgf10^K-v2^*^/+^; *tdTomato^flox/flox^* mice, indicating that the line is not leaky. By contrast, a solid signal was found in Tam-treated *Fgf10^Ki-v2^*^/+^; *tdTomato^flox/flox^* lungs. A weak signal was detected in Tam-treated *Fgf10^Ki-v1^*^/+^; *tdTomato^flox/flox^* lungs as described in a previously study [14] (Fig. 6B). Flow cytometry analysis was also conducted to quantify the total number of Tom^Pos^ cells in both conditions as well as their identity [14]. Only 0.3% Tom^Pos^ cells over total number of cells were detected in *Fgf10^Ki-v1^*^/+^; *tdTomato^flox/flox^.* This number is in line with the previously reported 0.1% [14] and confirms that the *Fgf10^Ki-v1^*^/+^ line is not efficient to target Fgf10^Pos^ cells in the adult lung. By contrast, we observed 5.8% of Tom^Pos^ cells over total cells in *Fgf10^Ki-v2^*^/+^ lungs. Further analysis showed that these cells were mostly Cd31^Neg^Cd45^Neg^Epcam^Neg^ cells (85 %) identifying them as resident mesenchymal cells (rMC). 14.3% of the Tom^Pos^ cells were also Sca1^High^, a functional marker of the rMC subpopulation capable of sustaining the self-renewal of AT2 stem cells in the alveolosphere organoid model [15]. Of note 53% of the Sca1^High^ were LipidTox^Pos^ identifying as lipofibroblasts (LIFs) Altogether these results suggest that TomPos cells are heterogeneous and comprise a significant percentile of LIFs as previously reported [14].

**Figure 6.**
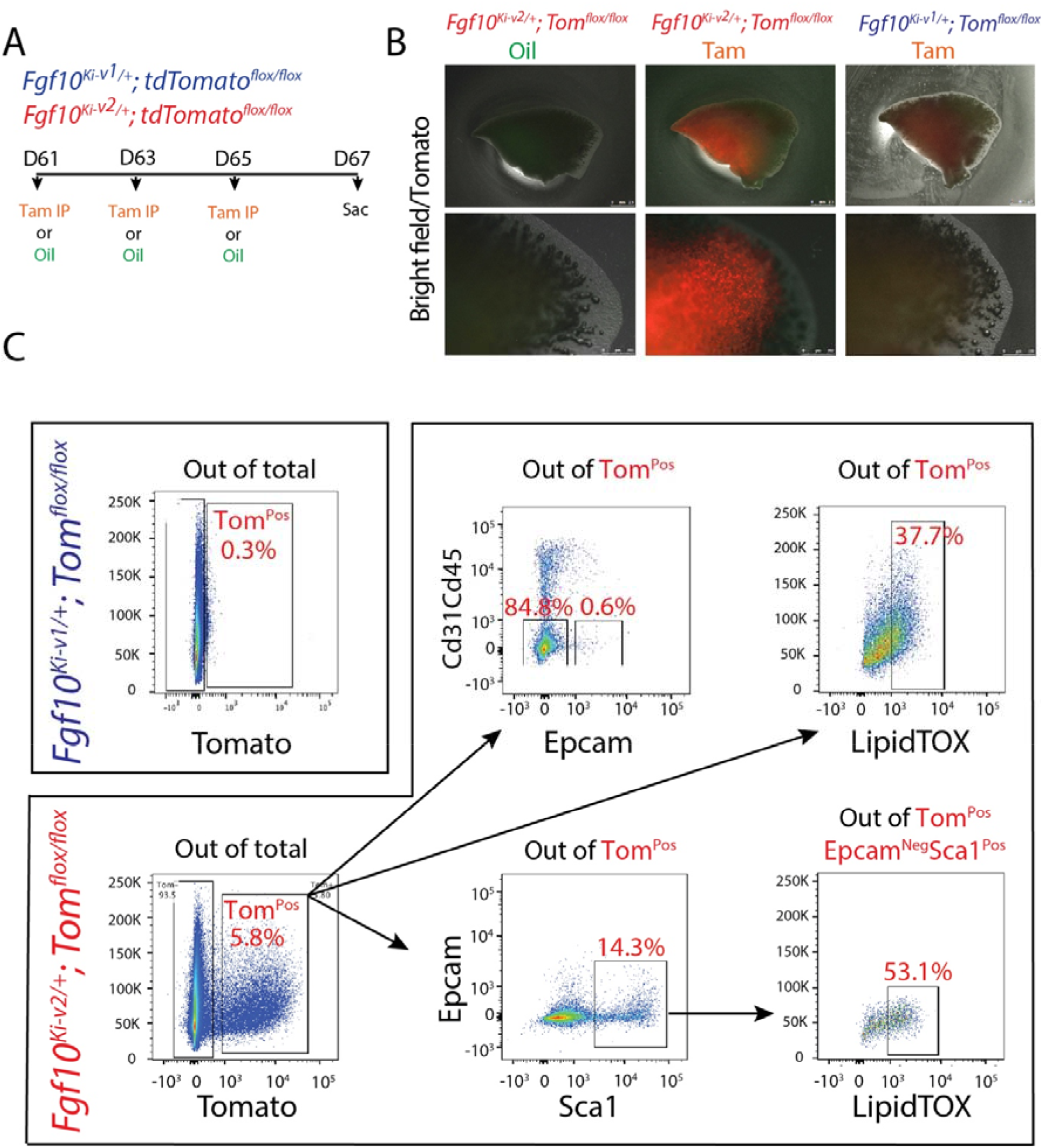
*Fgf10^Ki-v2/+^*; *tdTom^flox/+^* adult lungs display enhanced number of Tom^Pos^ cells compared to the *Fgf10^Ki-v1/+^*; *tdTom^flox/+^* adult lungs. **(A)** 2-months-old *Fgf10^+/+^*; *Fgf10^Ki-v1/+^*; *tdTom^flox/+^* and *Fgf10^Ki-v2/+^*; *tdTom^flox/+^* mice received 3 Tam IPs at P61, P63 and P65 and were sacrificed at P67. **(B)** Whole-mount fluorescent images of WT, *Fgf10^Ki-v2/+^*; *tdTom^flox/+^* and *Fgf10^Ki-v1/+^*; *tdTom^flox/+^* lungs at P67. Higher magnification of the lungs are shown in the lower panel. **(C)** Flow cytometry analysis of *Fgf10^Ki-v2/+^*; *tdTom^flox/flox^* mice lung homogenate.

### The 3’UTR region for *Fgf10* gene contains many key transcription factor binding sites

The decrease in *Fgf10* expression in *Fg10^Ki-v2^* mice suggested that important transcription factor binding sites (TFBS) were impacted by the genetic manipulation in the 3’UTR of the *Fgf10* gene. We determined the identity of TFBS located at proximity of the 3’UTR of the *Fgf10* gene using an online TFBS prediction tool. We compared these TFBS with previously published TF expressed in the lung mesenchyme [8]. We found several key TFBS matching TF expression such as *Hoxa5*, *Pou3f1*, *Pou2f1*, *Foxm1* and *Meis1* (Fig. 7). Mutant *Hoxa5* mice shows decreased surfactant production and disrupted tracheal cartilage, leading to respiratory distress and low survival rate at birth. *Pou3F1*, also known as *Oct6* is primarily expressed in neural cells. Mutation of *Pou3F1* also cause lethality at birth due to respiratory distress [16] [17, 18]. The body size of *Pou2f1* knockout embryos are smaller. Knock out of *Pou2f1* shows smaller body size of embryos and fully lethality at birth [19]. *Foxm1* expression is crucial to both epithelial and mesenchymal layer. It is found increased smooth muscles around proximal airways and reduced pulmonary microvasculature due to conditional inactivation of Foxm1 in lung mesenchyme [20]. In lung epithelium, the conditional inactivation of Foxm1 cause the reduction in sacculation and a delayed differentiation of alveolar type I cells. [21]. Knockout of *Meis1* caused fully lethality during embryonic stage around E14.5 due to microvascular and hematopoietic defects in lung [22]

**Figure 7.**
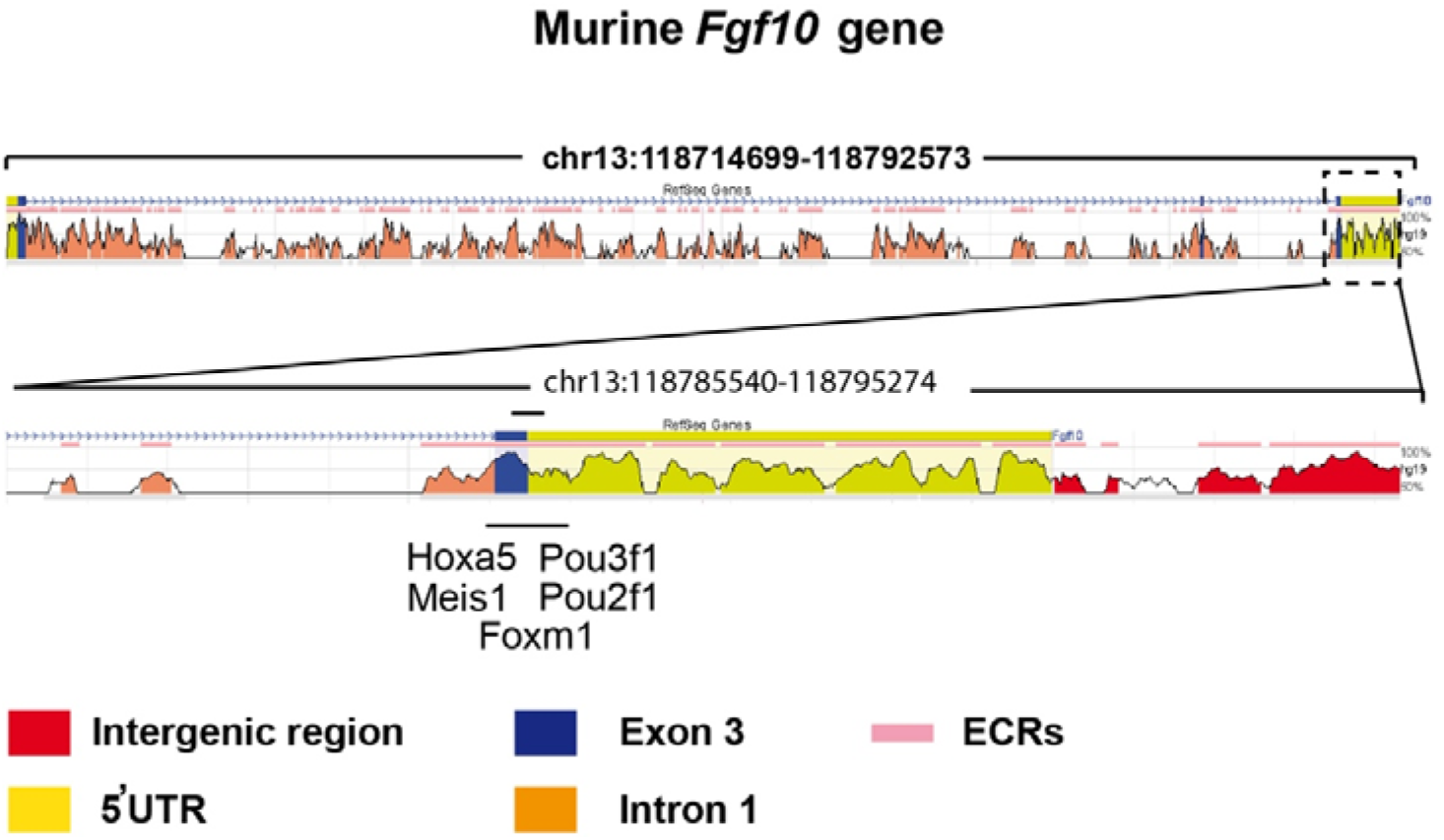
Bioinformatics analysis identified potential transcriptional factors binding sites in the 3’UTR of Exon 3. (A) The murine Fgf10 gene is made of 3 exons. (B) High magnification of Exon 3 and the associated 3’UTR. Several putative important transcriptional factors binding sites were found in this region.

## Discussion

Fgf10 is an essential morphogen underlying the developmental process of multiple organs including the lung. FGF10 signaling is also crucial during homeostasis and in the process of injury/repair in the adult lung. *FGF10* dysregulation in human has been implicated in some major respiratory diseases, such as bronchopulmonary dysplasia (BPD), Idiopathic pulmonary fibrosis (IPF) and chronic obstructive pulmonary disease (COPD). For example, Increased FGF10 expression level in IPF patients has been found [23]. However, FGF10 expression is inversely correlated to the disease progression with higher levels in stable IPF vs. lower level in end-stage IPF. Higher FGF10 expression in the early, stable stage of IPF is most likely correlated with the repair process. Insufficient FGF10 level in prematurely newborn infants is associated with arrested lung development at the saccular stage [24]. *Fgf10* deficiency in a newborn mouse model of hyperoxia-induced BPD led to drastic increase in lethality associated with abnormal alveolar epithelial type 2 (AT2) cell differentiation as well as surfactant production [25].

Fgf10 performs also a key function for the repair of the bronchial epithelium after injury [5]. Our knowledge about the sources of Fgf10 in this context has been evolving. FGF10 was first described to be expressed by airway smooth muscle cells (ASMCs) [5], whereas more recent work identified a peribronchiolar mesenchymal population capable of producing Fgf10 during the repair process, which is not derived from the ASMCs [4].

In COPD, the conducting airway epithelium undergoes massive remodeling causing an irreversible airway obstruction [26]. Interestingly, we have reported that the Fgf10-Hippo epithelial mesenchymal crosstalk also maintains and recruits lung basal stem cells in the conducting airways [27]. While transient *Fgf10* expression by ASMCs is critical for proper airway epithelial regeneration in response to injury, sustained Fgf10 secretion by the ASMC niche, in response to chronic Ilk/Hippo inactivation, results in pathological changes in airway architecture resembling the abnormalities seen in COPD. The inhibition of FGF10/FGFR2b signaling may therefore be an interesting approach to treat chronic obstructive airway lung diseases. Conversely, the opposite situation might occur in the respiratory airways in that destruction of the alveolar compartments resulting in emphysema may be due to insufficient FGF signaling. Interestingly, recombinant FGF7 has been reported to induce de novo-alveologenesis in the elastase model of emphysema in mice [28].

The previous *Fgf10^Ki-V1^* model was mainly used to trace the Fgf10^Pos^ cells during embryonic development. A near complete loss of the labeling capacity of Fgf10^Pos^ cells during postnatal stages limited its utilization in the analysis of their cell fate in adult lung homeostasis and during the process of injury/repair. In order to overcome the limitations of the *Fgf10^Ki-V1^* line, we generated and validated this new knock in *Fgf10^Ki-V2^* line. Upon crossing with a tdTomato reporter line, we demonstrated that the tdTomato expression domain faithfully reproduced the previously reported the *Fgf10* expression pattern [6], and a more robust labeling of Fgf10^Pos^ cells was achieved in the postnatal stages in spite of a mismatch between Cre and Fgf10 expression, which could be explained by the disruption of critical transcription factor binding sites located in the 3’UTR of the *Fgf10* gene. Therefore, this line will be a valuable tool to define further the mesenchymal cell populations in the adult lung contributing to the repair process after injury. Combined crosses with existing or novel Dre-ERT2 recombinase driver lines may allow to capture subpopulations of Fgf10^Pos^ cells/lineages based on the expression of two markers [29]. The main Fgf10^Pos^ subpopulation is represented by the lipid-containing alveolar interstitial fibroblasts (lipofibroblasts or LIFs). LIFs are increasingly recognized as an important component of the AT2 stem cell niche in the rodent lung. Although LIFs were initially believed merely to assist AT2 cells in surfactant production during neonatal life, recent evidence suggests that these cells are indispensable for self-renewal and differentiation of AT2 stem cells during adulthood [7]. Despite increasing interest in lipofibroblast biology, little is known about their cellular origin or the molecular pathways controlling their formation during embryonic development. We have shown that a population of LIF progenitors emerges in the developing mouse lung between E15.5 and E16.5 and that Fgf10 is critical for their differentiation towards the LIF lineage [30]. We have also reported the existence of Fgf10^Pos^-LIF as well as Fgf10^Neg^-LIFs [30]. The difference between these two populations is still unclear and will require further studies. In the context of bleomycin-induced lung fibrosis, in vivo lineage tracing indicate that LIFs transdifferentiate into activated myofibroblast during fibrosis formation and that a significant proportion of the labeled activated myofibroblasts transdifferentiate back to LIFs during fibrosis resolution [31].

In conclusion, we have successfully generated a new *Fgf10^CreERT2^* line with enhanced labeling efficiency of Fgf10^Pos^ cells postnatally. This line, which displays normal expression of *Fgf10* in *Fgf10^CreERT2/+^*, avoids many developmental defects linked to deficient *Fgf10* expression. Therefore, It paves the way for performing cell-autonomous based studies to investigate the role of these Fgf10^Pos^ cells as well as associated signaling pathways during lung development and disease.

## Acknowledgements

SB was supported by the Cardio-Pulmonary Institute and by grants from the Deutsche Forschungsgemeinschaft (DFG; BE4443/1-1, BE4443/4-1, BE4443/6-1, KFO309 P7, and SFB1213-projects A02 and A04). S.B. also acknowledges the support of the CPI. CC was supported by the Interventional Pulmonary Key Laboratory of Zhejiang Province, the Interventional Pulmonology Key Laboratory of Wenzhou City, the Interventional Pulmonology Innovation Subject of Zhejiang Province, the National Nature Science Foundation of China (81570075, 81770074), Zhejiang Provincial Natural Science Foundation (LZ15H010001), Zhejiang Provincial Science Technology Department Foundation (2015103253), the National Key Research and Development Program of China (2016YFC1304000).

---

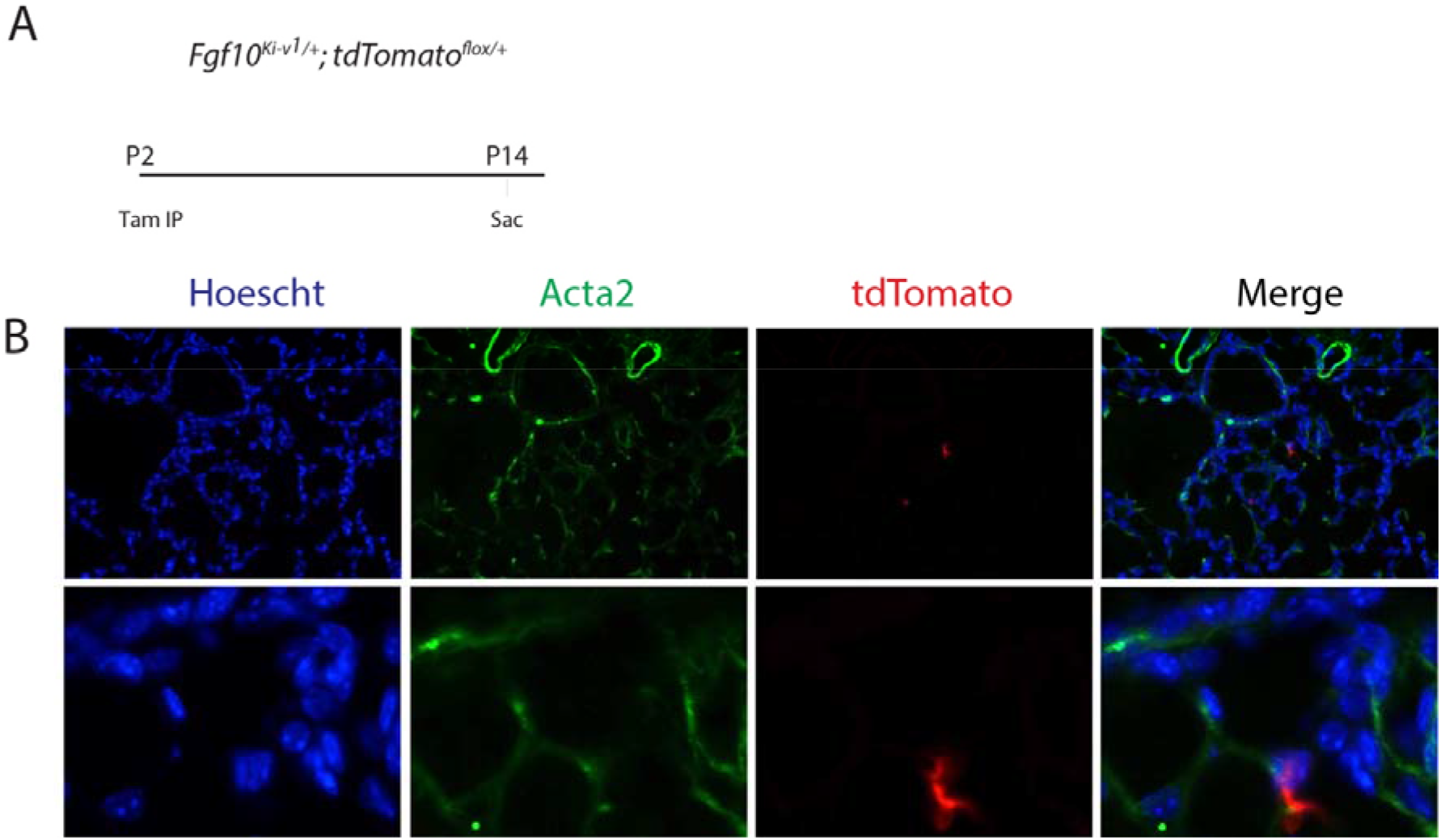

